# ProteinDJ: a high-performance and modular protein design pipeline

**DOI:** 10.1101/2025.09.24.678028

**Authors:** Dylan Silke, Julie Iskander, Junqi Pan, Andrew P. Thompson, Anthony T. Papenfuss, Isabelle S. Lucet, Joshua M. Hardy

**Author notes:** Co-corresponding authors: Isabelle S. Lucet, Joshua M. Hardy.

## Abstract

Leveraging artificial intelligence and deep learning to generate proteins *de novo* (a.k.a. ‘synthetic proteins’) has unlocked new frontiers of protein design. Deep learning models trained on protein structures can generate novel protein designs that explore structural landscapes unseen by evolution. This approach enables the development of bespoke binders that target specific proteins and domains through new protein-protein interactions. However, successful binder generation can suffer from low *in silico* success rates, often requiring thousands of designs and hundreds of GPU hours to obtain enough hits for experimental testing. While workstation implementations are available for binder design, these are limited in both scalability and throughput. There is a lack of efficient open-source protein design pipelines for high-performance computing (HPC) systems that can maximise hardware resources and parallelise the workflow efficiently.

Here, we present ‘ProteinDJ’—an implementation of a synthetic protein design workflow that is deployable on HPC systems using the Nextflow portable workflow management system and Apptainer containerisation. It parallelises the workload across both GPUs and CPUs, facilitating generation and testing of hundreds of designs per hour, accelerating the discovery process. ProteinDJ is designed to be modular and includes RoseTTAFold Diffusion (RFdiffusion) or BindCraft for fold generation, ProteinMPNN or Full-Atom MPNN (FAMPNN) for sequence design, and AlphaFold2 or Boltz-2 for prediction and validation of designs and binder-target interfaces, with supporting packages for structural evaluation of designs. ProteinDJ democratises protein binder design through its robust and user-friendly implementation and provides a framework for future protein design pipelines. ProteinDJ is freely available at https://github.com/PapenfussLab/proteindj.

## Introduction

Proteins are fundamental biological molecules that perform a wide range of functions in nature, including enzymes that catalyse chemical reactions, transcription factors that regulate gene expression, and antibodies that recognise foreign pathogens. These functions can be regulated through protein-protein interactions (PPIs)^1^. Consequently, the production of novel PPI binders that can specifically activate or inhibit protein function is of immense biomedical and therapeutic interest^2^. This is exemplified by the joint award of the 2024 Nobel Prize in Chemistry to David Baker, Demis Hassabis, and John Jumper for computational protein design and protein structure prediction^3^.

Recent advances in artificial intelligence, particularly deep learning approaches, have demonstrated remarkable ability to navigate the sparse solution space of protein binder design by learning underlying patterns and relationships from known protein structures^4; 5^. The current state-of-the-art process for *de novo* protein binder generation includes (1) RFdiffusion^5^ or BindCraft^6^ for structure generation, (2) ProteinMPNN^7^ for sequence assignment and (3) AlphaFold2 Initial Guess^4; 8^ for filtering and interaction prediction. Though newer methods like AlphaProteo^9^, BoltzDesign1^10^ and BoltzGen^11^ demonstrate promising results, RFdiffusion and BindCraft are the most validated benchmarks for structure generation. Similarly, alternate software for filtering and interaction prediction exists, including AlphaFold3, Boltz-2 and Chai-1^12–14^, and new structure prediction metrics have emerged for filtering and ranking designs^15–17^. The rapidly advancing nature of this field necessitates versatile and modular workflows to evaluate and implement continuing developments.

Structure generation by RFdiffusion is a foundational technology for protein design. It employs a diffusion model – a deep learning model that learns to progressively build plausible protein structures from Gaussian noise^5^. For binder generation, this construction process is guided by the target protein surface to create complementary interfaces. It has demonstrated success in generating binders with nanomolar affinity for diverse therapeutic targets, including binders for pathogenic proteins (e.g. influenza hemagglutinin, snake venom) and human cell surface receptors (e.g. insulin receptor, programmed death-ligand 1, platelet-derived growth factor receptor α)^5; 18; 19^.

More recently, BindCraft has emerged as a powerful alternative binder design algorithm^6^. Instead of using diffusion, it uses backpropagation through the AlphaFold2 network to hallucinate new binders. In contrast to RFdiffusion, which keeps the target residues fixed during binder design, BindCraft re-predicts the interface at each interaction and facilitates development of ‘induced-fit’ target-binder interfaces. Despite the significantly increased runtime of the iterative hallucination process compared to RFdiffusion, BindCraft was able to achieve a similar numbers of successful designs per GPU hour^6^. As with RFdiffusion, BindCraft designs are passed to ProteinMPNN for sequence optimisation and evaluation is performed using AlphaFold2 structure prediction. BindCraft was able to generate successful binders for 12 diverse targets including extracellular receptors and intracellular enzymes^6^, and was recently used to create the highest affinity binder to epidermal growth factor receptor in an international crowdsourced binder design competition^20^.

The success of synthetic binder generation can vary greatly depending on the target protein interface. Achieving enough designs that pass *in silico* metrics and succeed in experimental testing can require thousands of designs and hundreds of GPU hours. While workstation implementations are available for binder design, these are limited in both scalability and throughput required for the broad application of protein binder design. To meet these needs, *de novo* protein design pipelines must be adapted to HPC systems that can maximise utilisation of hardware resources and parallelise the workflow efficiently. For research institutes with several research groups designing binders against multiple targets, scalable workflows are crucial in preventing bottlenecks and delivering timely results. Currently, there is a lack of efficient implementations of such pipelines. Additionally, deployment of these computational tools presents significant technical barriers, requiring careful management of various CUDA versions and Python environments, which limits their accessibility to non-expert users. Finally, metadata from these tools often cannot be easily parsed and analysed using common data analysis tools like Microsoft Excel or pandas, which impedes efficient data processing workflows.

Motivated by these limitations, we developed ProteinDJ (https://github.com/PapenfussLab/proteindj) — a software package that utilises Nextflow^21^, a scientific workflow framework, to parallelize protein design tools for use with high-performance computing systems. It automates dependency management via Apptainer^22^ containers that contain pre-packaged applications to reduce setup time and efficiently distributes computation across both CPUs and GPUs. ProteinDJ introduces several key improvements to the standard pipeline with guidance for different types of protein designs. ProteinDJ includes a subprogram ‘Bindsweeper’ for parameter sweeping, allowing users to systematically explore different settings and identify optimal parameter combinations. Structure-guided filtering enables rejection of undesired folds early in the pipeline, reducing downstream computational costs. To facilitate analysis, ProteinDJ also provides improved design metrics and summary statistics for the filtered outputs. Ultimately, this pipeline aims to improve accessibility and performance of protein binder design and enable uptake of these powerful yet complex software tools.

## Results

### ProteinDJ is a high-performance and parallelisable protein design pipeline

The ProteinDJ pipeline is divided into four major stages: (1) fold design, (2) sequence design, (3) structure prediction, and (4) analysis and reporting (Figure 1). It integrates different software packages with options for sequence design and structure prediction (Supplementary Table 1). For fold design, we have integrated RFdiffusion and BindCraft, which use diffusion and hallucination methods respectively to design protein folds. The user can choose from sequence design with ProteinMPNN or the more recent tool FAMPNN that extends ProteinMPNN to iteratively fit side-chains during sequence design^23^. There is only one set of weights available for Full-atom MPNN, but ProteinMPNN can access three different models: ‘vanilla’, ‘soluble’ (a.k.a. SolMPNN)^24^, and ‘hyper’ (a.k.a. HyperMPNN)^25^. ProteinMPNN can also optionally perform an energy minimisation of the binder structure (using the Rosetta FastRelax protocol)^4^. This is a CPU intensive and relatively slow step but allows for iterative sequence design with cycles of ProteinMPNN and Rosetta FastRelax. For structure prediction, there is a choice of Boltz-2^12^ or a modified form of AlphaFold2 that preserves the target structure while predicting the binder structure and position (known as AlphaFold2 Initial Guess)^4^, both with GPU acceleration.

**Figure 1.**
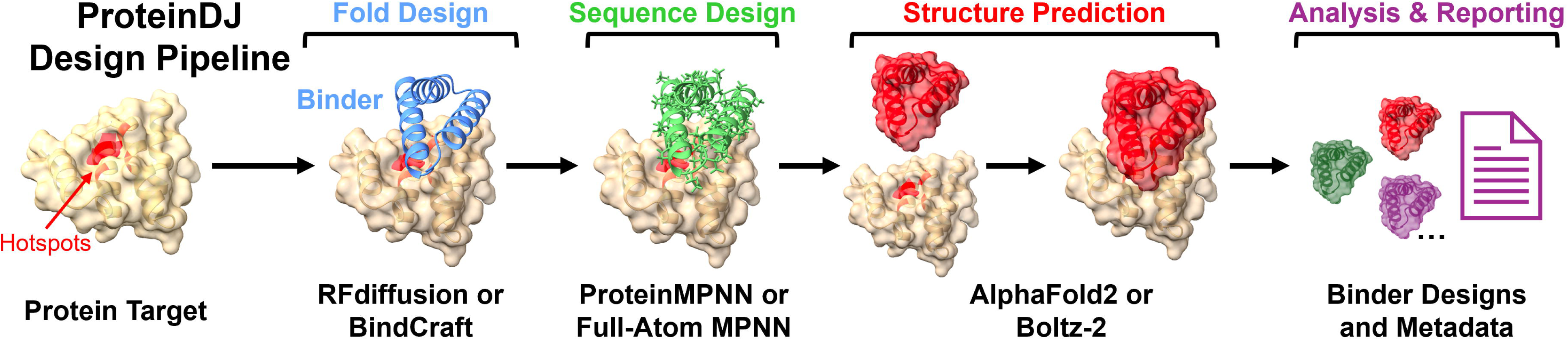
The four major stages of the ProteinDJ pipeline for *de novo* binder design. RFdiffusion or BindCraft is used to design folds near hotspots on a protein target. The user can choose from ProteinMPNN or Full-Atom MPNN for sequence design, and AlphaFold2 Initial Guess or Boltz-2 for structure prediction and validation.

We designed the pipeline to parallelize stages across available GPUs and CPUs using Nextflow and HPC schedulers (e.g. SLURM), with synchronization points across GPU to CPU stage transitions. ProteinDJ is highly configurable via a centralized Nextflow configuration file enabling users to fine-tune each aspect of the design pipeline. A comprehensive overview of each configuration parameter, their descriptions, and default values is provided in the Methods.

The workflow begins by passing configuration parameters and any input files (Figure 2a) to RFdiffusion or BindCraft to generate new secondary and tertiary structures (Figure 2b). Key configuration parameters include the number of requested designs, the choice of sequence design or structure prediction software, and the output directory (see Methods). The total number of requested designs is divided into batches per GPU to maximise parallel throughput. Generated folds then transition to a filtering stage, where the radius of gyration (compactness) and the number of alpha-helices and beta-strands are counted, allowing rejection of low complexity folds (e.g. single alpha-helices) and compact or elongated binders depending on the need of the design case. Folds meeting these criteria then proceed to either ProteinMPNN (with or without FastRelax) or FAMPNN for sequence assignment of the RFdiffusion poly-glycine binders or refinement of the BindCraft sequences (Figure 2c). The user determines the number of sequences per fold to be generated, and the models are distributed across available CPUs for ProteinMPNN or GPUs for FAMPNN. The outputs of the sequence generation step move to a second filtering stage where designs with poor sequence scores are rejected. Examples of criteria for rejection include negative log-likelihood for ProteinMPNN or predicted sidechain error for FAMPNN.

**Figure 2.**
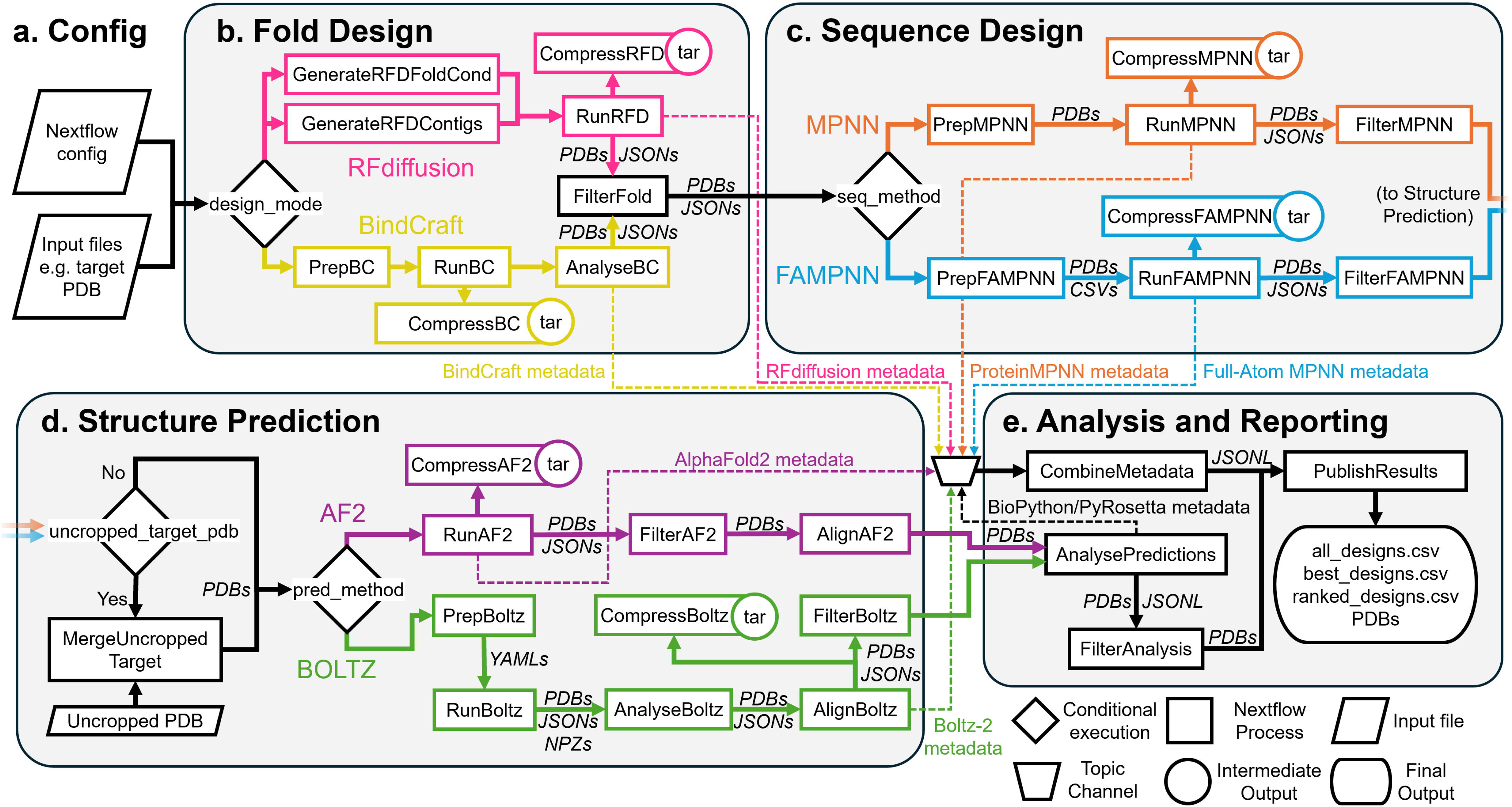
Data flow and connectivity diagram of ProteinDJ. **a.** User parameters and files are provided to Nextflow to execute a ProteinDJ run. **b.** In the first stage, the user selects a ‘design_mode’ that will either launch RFdiffusion or BindCraft to generate folds (RunRFD/RunBC) with preparatory processes depending on the mode selected (GenerateRFDcontigs/GenerateRFDFoldCond/PrepBC). All output PDBs and metadata (JSON files) are captured and compressed as intermediate results (CompressRFD/CompressBC). BindCraft outputs are analysed and transformed for downstream compatibility (AnalyseBC) and passed to a filtering stage (FilterFold) that also handles RFdiffusion outputs. Metadata from these processes are exported to a shared ‘topic’ channel for metadata collection and integration. **c.** In the sequence design stage, the user can select between two packages using the ‘seq_method’ parameter. Each has a preparatory stage (PrepMPNN/PrepFAMPNN) in which metadata from Fold Design are used to mask residues during sequence design (for RunMPNN and RunFAMPNN respectively). Similarly to RFdiffusion, intermediate outputs are captured and compressed (CompressMPNN/CompressFAMPNN), before filtering (FilterMPNN/FilterFAMPNN). **d.** The user can optionally provide an uncropped target PDB file for structure prediction that is merged with designs in MergeUncroppedTarget. There are two methods available for structure prediction (RunAF2 and RunBoltz), selected using the ‘pred_method’ parameter. When selecting Boltz-2 for prediction, the PrepBoltz stage is used to generate a YAML containing the sequences of the designs and AnalyseBoltz calculates additional structure metrics for the predictions. For ease of visualisation, the designs are aligned to each other (AlignAF2/AlignBoltz), and a third filtering step is optionally performed (FilterAF2/FilterBoltz). **e.** In the analysis and reporting stage, the remaining PDB files are subjected to further analysis (AnalysePredictions) with a final filtering stage (FilterAnalysis), and these metrics are combined with other metrics from other stages into a CombineMetadata process. This generates multiple CSV files, one with metadata of all designs (all designs.csv), designs that passed filtering if enabled (best_designs.csv), and optionally designs ranked by prediction metrics (ranked_designs.csv). The CSV files are published along with the PDB files that survived filtering (PublishResults).

In the next stage of the pipeline, sequence designs are validated by structure prediction using GPU-parallelized Boltz-2 or AlphaFold2 Initial Guess (Figure 2d). As RFdiffusion/BindCraft runtime scales exponentially with target size for binder design tasks, it is common practice to truncate input target structures. While this improves runtime, it can inadvertently lead to the design of binders that clash with truncated components of the target structure or that interact with exposed hydrophobic residues. To address this, we have designed and implemented optional functionality that replaces the target chain in the design PDB file with an uncropped target structure provided by the user before structure prediction. A structure prediction filtering stage is used to filter out designs with low prediction confidence (e.g. predicted aligned error) or with significant differences to the input fold.

Designs that pass structure prediction filters are aligned and proceed to a final analysis stage (Figure 2e), that uses PyRosetta^26^ and Biopython^27^ to generate additional metrics, as described in Supplementary Table 3. These metrics can be used to optionally filter out designs with poor biophysical properties e.g. hydrophobicity, in a final filtering process. Metrics from the entire pipeline are subsequently collated into a CSV file for user perusal. The designs that passed filtering are output to the results directory as aligned PDB files for easy comparison and visualisation (e.g. in ChimeraX^28^) and can be optionally ranked to select the top designs according to metrics correlated with experimental binding (e.g. pae_interaction for AlphaFold2, and ipSAE_min for Boltz)^29^. A summary of the workflow and success rates is printed to the log and output (Supplementary Figure 1).

Despite the complexity of the workflow, ProteinDJ exhibits scaled efficiently across multiple GPUs, achieving near-linear reductions in runtime compared to the serial baseline, with minimal loss in parallel efficiency. Using 8 A30 GPUs, ProteinDJ generated and evaluated 500 binder folds and 4000 unique sequences using ProteinMPNN and AlphaFold2 in a run time of 2 hours and 26 minutes, compared to 16 hours and 55 minutes when using a single GPU (See Methods and Supplementary Figure 2a). We observed a parallel efficiency of 97.0%, 93.6%, and 86.5% over 2, 4, and 8 NVIDA A30 GPUs respectively (Supplementary Figure 2b).

We also compared the efficiency of each software package in generating 1,000 binders (100 folds, 10 sequences per fold) against a benchmarking target, Programmed cell death 1 ligand 1 (PD-L1) (Supplementary File 1, Supplementary Table 6). RFdiffusion generated a fold every 54.9 seconds on average compared to 620.5 seconds per fold with BindCraft. ProteinMPNN designed a sequence every 1.3 seconds and slowed to 56.9 seconds per design when performing one FastRelax cycle, whereas FAMPNN required only 4.8 seconds per sequence. Boltz-2 was more efficient than AlphaFold2 Initial Guess and generated a prediction every 8.6 seconds on average compared to an average of 16.4 seconds per prediction with AlphaFold2 Initial Guess.

### Modular design enables future software integrations

To ensure modularity of the workflow and facilitate integration of new software, we standardised the data and metadata formats. Each design is passed as a single PDB file, and rather than embed the metadata into the PDB file, which can be difficult to extract later, we embed all metadata from the stage into a JSON file. To pair designs with metadata, we used a unique fold number (fold_id), to differentiate each RFdiffusion/BindCraft design, and a unique sequence number (seq_id) once a sequence has been generated for a fold. Each metadata filename and entry contain these ID fields so that metadata can be merged at the end of the pipeline into a user-friendly CSV format. The IDs also allow batching of each PDB and JSON file pair into different batch sizes as appropriate for CPU or GPU processing.

Rather than pass each metadata file to the next software package, we collect the metadata using a Nextflow topic channel that can consume outputs from different processes. This ensures metadata is published at each stage of the workflow and captured even if the pipeline is interrupted. By only passing the PDB files between stages, we decoupled metadata processing between different software packages, allowing for alternative workflows and software options. Although compressed and storage efficient formats exist for PDB files (e.g. the ‘silent’ format from https://github.com/bcov77/silent_tools), we did not implement them as it complicates batching of PDB files in Nextflow. To minimize impact on storage, however, we compress PDB and metadata files before publishing to the output directory.

### Implementation of modes to RFdiffusion delineate and simplify protein design workflows

RFdiffusion is a flexible protein design software that contains a plethora of optional parameters. To increase accessibility of the pipeline to new users, we have divided the pipeline into monomer and binder design workflows with four modes for each: ‘*de novo*’, ‘fold conditioning’, ‘partial diffusion’ and ‘motif scaffolding’ (Figure 3). These modes organise the existing optional parameters of RFdiffusion into defined applications where they are relevant. By streamlining the pipeline entry-points this way, we aim to reduce unexpected conflicts and errors during execution from incompatible RFdiffusion parameters (Supplementary Table 4).

**Figure 3.**
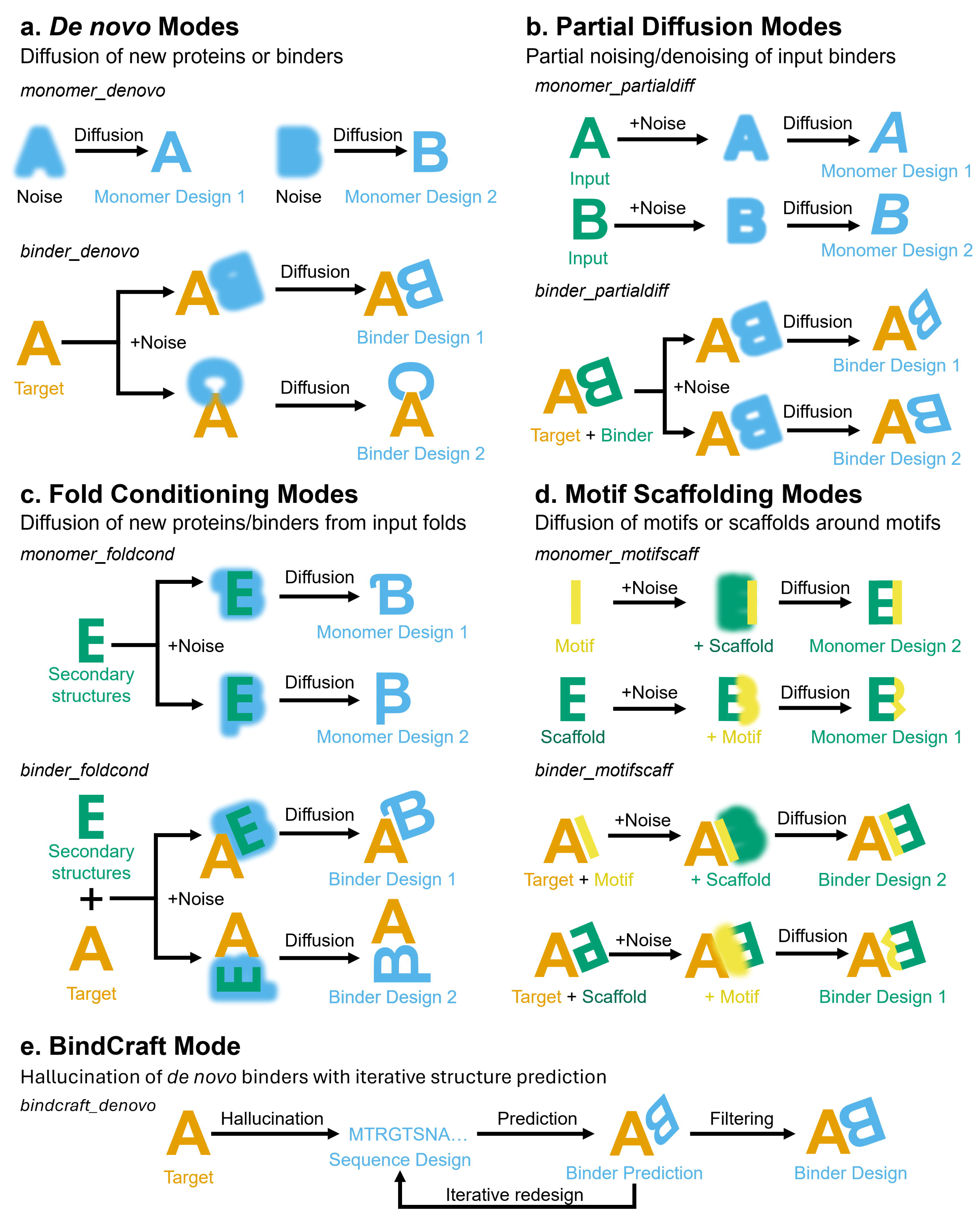
Protein design modes for ProteinDJ. **a.** *De novo* protein design is used to diffuse new proteins from noise, either in isolation (‘monomer_denovo’) or with an input chain (‘binder_denovo’). **b.** Partial diffusion can be used to generate variations of input proteins (‘monomer_partialdiff’) or binders (‘binder_partialdiff’) by noising and denoising structures. **c.** Fold conditioning is a constrained form of *de novo* protein design in which the designs must follow secondary structure definitions provided during inference and can be used for monomers (monomer_foldcond) or binders (binder_foldcond). **d.** Motif scaffolding provides the ability to inpaint or outpaint existing structures, while preserving key regions of the input structure. **e.** BindCraft runs within a separate design mode for *de novo* binder design, utilising iterative hallucination instead of diffusion with internal filtering of designs.

The simplest *de novo* mode, ‘monomer_denovo’, diffuses a single chain in each design and requires only specification of the chain length. The ‘binder_denovo’ mode diffuses binders to a target protein derived from an input PDB file with the option to provide specific residues, called ‘hotspots’, to focus the design on (Figure 3a). This approach can be used to generate new binders to targets that lack biological adaptors or existing binders. The partial diffusion modes, ‘monomer_partialdiff’ and ‘binder_partialdiff’, will noise and denoise an input monomer or binder (or parts thereof) to generate variants, according to the number of diffusion timesteps provided (Figure 3b). Partial diffusion can be used to optimise candidate binders from *de novo* methods.

The fold conditioning modes, ‘monomer_foldcond’ and ‘binder_foldcond’, allow users to guide diffusion of new proteins with specific folds (Figure 3c). This requires generation of files containing secondary structure information from a library of scaffold structures. To assist users in binder design, we have generated scaffold libraries of four fold types using PDB files from a previously published minibinder study^30^: 3-helical and 4-helical bundles, and two mixed alpha-helical and beta-stranded folds based on ferredoxin and a synthetic fold (Figure 4). Although this mode lacks the variability of designs produced by *de novo* modes, the folds are more likely to be recapitulated during structure prediction and this increases the *in silico* success rate (as previously observed^5^). RFdiffusion also has a strong bias towards alpha-helical folds when performing *de novo* fold design that is somewhat mitigated by a ‘beta’ checkpoint model for RFdiffusion^5^, but fold conditioning allows for explicit use of beta-sheet rich structures that may be preferred in certain design contexts.

**Figure 4.**
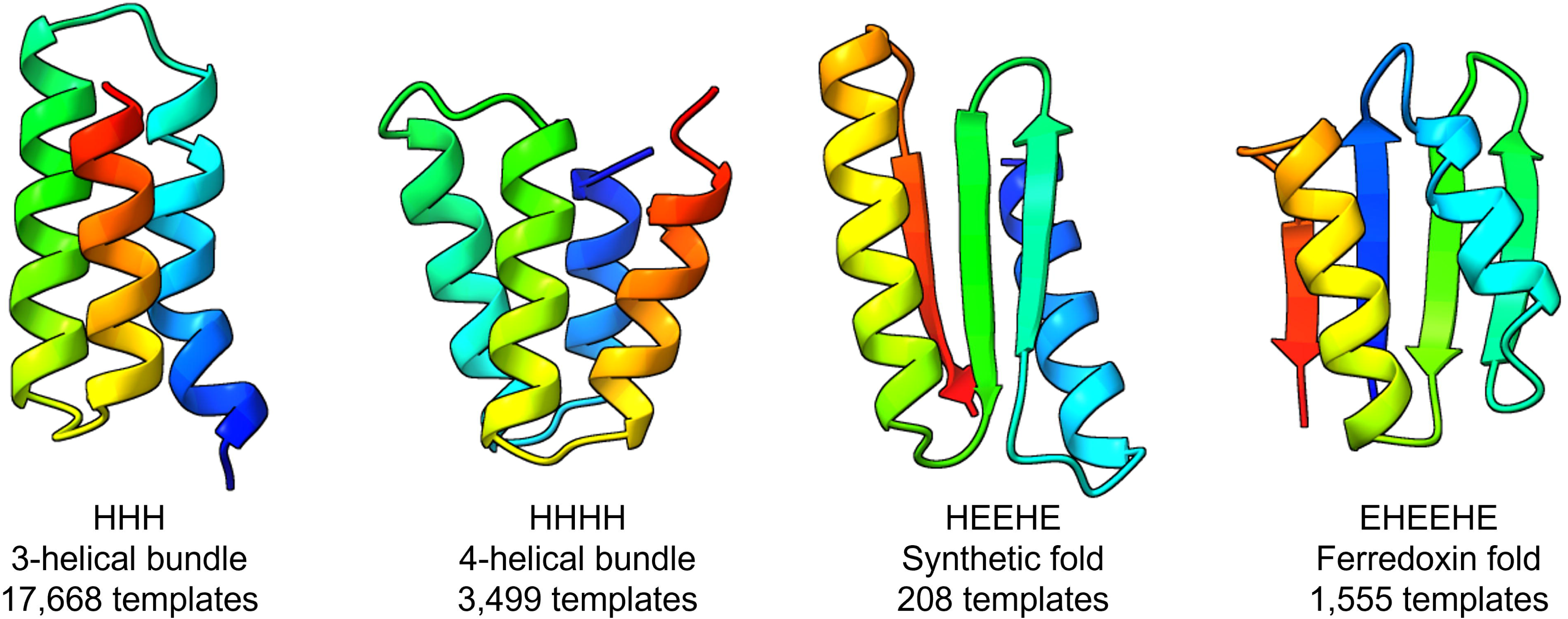
Examples of the four scaffold types provided for binder fold conditioning containing alpha-helical (H) and beta-strand (E) secondary structures. Structures are rendered as cartoons with rainbow colouring from N- to C- termini in ChimeraX^28^.

The motif scaffolding modes, ‘monomer_motifscaff’ and ‘binder_motifscaff’, are the most complex and require specification of specific residues or regions for diffusion (Figure 3d). These modes can be used to modify regions of input proteins while preserving the rest of the structure and/or sequence. This can be used to diffuse a scaffold around a binding motif, or to diffuse a binding motif into a scaffold. To facilitate this, we directly map the RFdiffusion sequence masks to ProteinMPNN/FAMPNN to allow for partial changes to design sequences (see Methods).

BindCraft has its own mode, ‘bindcraft_denovo’, that is capable of performing *de novo* binder design and uses the same input parameters as RFdiffusion (e.g. hotspot_residues, input_PDB) with optional parameters specific to BindCraft (Figure 3e). The user can choose from different protocols for binder design e.g. ‘default’ vs ‘betasheet’, and whether to preserve the interface residues of the binder designed by BindCraft during the subsequent sequence design stage.

Our implementation of modes allow for differentiation in the downstream analysis of designs. For example, in monomer design options, the metrics can simply be calculated for all residues because there is only one chain, whereas in binder modes, metrics can be calculated for residues from the binder chain only and ignore target residues if applicable. The modes also facilitate the use of downstream software that have strict input requirements, such as a single target chain and binder chain only.

Some parameters are common to each mode, such as the number of designs and the output directory, but most optional parameters are restricted to specific modes (see Supplementary Table 4). For example, when performing partial diffusion of binders, the binder position is already known and thus specifying ‘hotspots’ is not meaningful, so this parameter is ignored. Likewise, providing ‘timesteps’ for partial diffusion is only relevant when a structure is provided and it is therefore disabled for *de novo* design. The specification of input residues and residues to diffuse, referred to as ‘contigs’ for RFdiffusion, can be an error-prone parameter for new users. We provide automatic generation of contigs for de novo and partial diffusion modes to include all residues from input PDBs. Experience users can specify contigs and this parameter is required for motif scaffolding modes. We have created schema files for running on the Nextflow Seqera Platform^21^ that show only relevant parameters for each mode with input validation and format checks during job configuration. We have also implemented input validation of files and syntax before launching RFdiffusion or BindCraft, for example, checking that the hotspot residue numbers are within the residue ranges provided for the input protein, a common error for new users.

### Development of a parameter sweeping algorithm for binder design - Bindsweeper

The most efficient parameters for binder design can vary between target proteins and identifying the ideal parameters can be laborious, requiring manual exploration of key parameters such as target hotspots. To enable users to automatically and systematically explore design parameters, we developed Bindsweeper – a Python-based command-line tool that can coordinate multiple ProteinDJ runs. Bindsweeper processes ProteinDJ parameters via command-line inputs and a YAML configuration file (Figure 5a), providing flexible execution options including a quick testing mode that tests each parameter combination with a smaller set of designs and parallelisation options.

**Figure 5.**
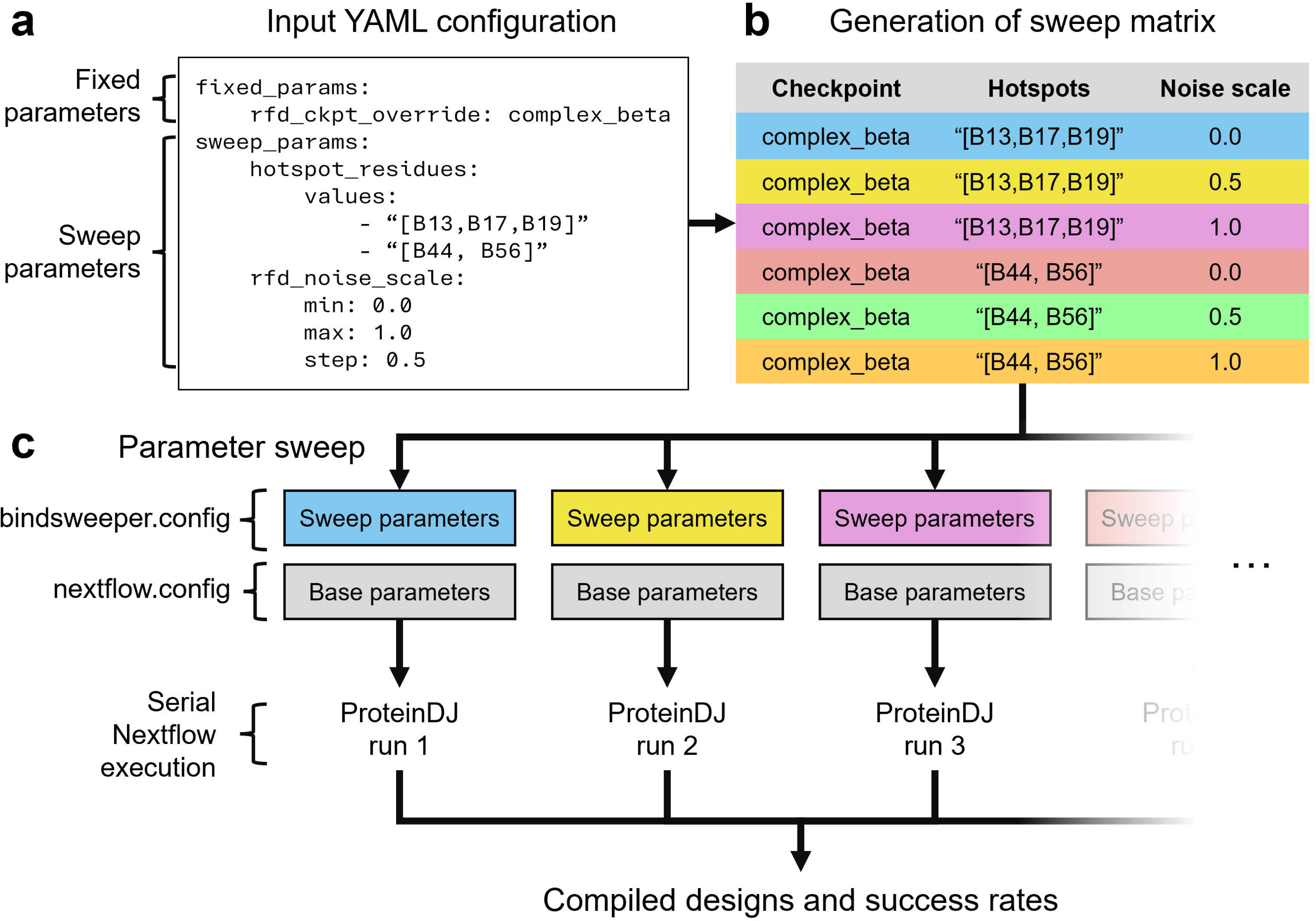
Workflow diagram for Bindsweeper parameter sweep algorithm. Bindsweeper uses an input YAML file (a) to generate a multi-dimensional matrix of sweep parameters (b), that can be used to perform sequential runs of ProteinDJ (c).

Bindsweeper generates a multi-dimensional parameter matrix by computing all possible combinations of specified sweep parameters (Figure 5b), supporting both list sweeps for discrete values and range sweeps for continuous values defined by start, stop, and step parameters. During execution (Figure 5c), Bindsweeper creates Nextflow profiles for each parameter combination, and executes pipeline runs with detailed timing and success tracking. After completing all parameter combinations, the results processor extracts putative binder PDB files, merges CSV outputs from each parameter combination, and performs success analysis to identify best-performing parameter combinations, with designs organized for downstream analysis and validation.

### Bindsweeper facilitates benchmarking and comparison of protein design software

To demonstrate the utility of Bindsweeper, we used it to benchmark and compare sequence design software. We used RFdiffusion to generate 100 *de novo* binder designs against each of the five benchmarking target proteins used in the RFdiffusion publication^5^ (Supplementary Table 6) and provided these to ProteinMPNN to generate 8 sequences for each, with and without a cycle of FastRelax, and with the standard ProteinMPNN model (a.k.a. ‘vanilla’ model) and the SolMPNN model. The same folds were also provided to FAMPNN and sequences were designed with and without fixing the position of target side-chains during design. After sequence design, structure prediction with AlphaFold2 Initial Guess was used to determine success rates using previously published metrics and thresholds (see Methods). The success rates varied dramatically between target proteins, in line with previous reports^5^. However, Bindsweeper revealed that the sequence design method also resulted in significant differences across the targets. With ProteinMPNN, one cycle of FastRelax improved the success rates for all targets, and the SolMPNN model outperformed the vanilla model for 3 of 5 targets. FAMPNN had the lowest success rates for 2 targets but performed better or similarly to ProteinMPNN in others. For most targets, fixing the side-chains on the target during sequence design with FAMPNN lowered the success rate.

## Discussion

In this work, we present a robust Nextflow pipeline for protein design that brings together diverse software packages in a unified and efficient framework. The pipeline incorporates several features that simplify and streamline protein design workflows. It scales with 86.5% efficiency across 8 GPUs, reducing the time to generate designs by almost an order of magnitude. Automated and detailed metadata management and reporting have been implemented to facilitate modularity and user control. Our uncropped structure prediction functionality reintroduces biological and structural context to the design process. Finally, we have delineated the software into simplified protein design tasks, with automated parameter selection, tailored to the user’s expertise. The software is supported by detailed documentation and examples of different design tasks.

ProteinDJ implements filtering steps after each design stage, which can be adjusted depending on design objectives, allowing the pipeline to be adapted for different protein design challenges. The filters also automatically reject designs without manual curation, saving valuable time. This is particularly valuable when targeting different protein-protein interaction types or when exploring novel fold spaces where standard filtering criteria may need adjustment. We provide metadata for rejected designs in one of the output CSV files (all_designs.csv), allowing users to see the scores before they were rejected and adjust the filters if needed to include. Additionally, we added new metadata on the biophysical and sequence characteristics of designs, such as the extinction co-efficient (i.e. whether the protein is detectable by UV spectroscopy) and the isoelectric point (pI) and molecular weight of the sequence, to support subsequent *in vitro* expression, purification and validation.

Our new tool Bindsweeper gives users the ability to automatically sweep parameter combinations for binder design. This enables systematic small-scale optimisation of parameters for target proteins such as the identification of hotspot residues, before embarking on large-scale design campaigns. This is especially important when success rates are so low as to prevent practical generation of sufficient binders for testing. Researchers must carefully balance the time spent on computational design prior to the more costly experimental validation process.

Bindsweeper also facilitates comparison and benchmarking of software packages within ProteinDJ, as we have demonstrated above. For example, we observed higher in silico success rates when using FastRelax in combination with ProteinMPNN compared to using ProteinMPNN alone, with the same input folds (Table 1). This improvement to success rates with RFdiffusion folds was similar to that observed by Watson et al.^5^, but while they reported inconsistent improvement to success rates across benchmarking targets, we observed a consistent improvement for the same targets. This could be due to differences in our benchmarking setups and whether the same input folds were used for each run. Whilst FastRelax demonstrates higher success rate, it does at the cost of increased run times per design. Further experimental studies are needed to evaluate whether this enhances success rates in vitro.

**Table 1.** Comparison of relative success rates of sequence inference methods for *de novo* binder design against each target. For each target, the same 100 folds were provided to each sequence design combination to generate 8 sequences each (n = 800). The highest success rate using AlphaFold Initial Guess for each target is bolded. HA = Influenza A H1 haemagglutinin, IL-7Rα = Interleukin-7 receptor alpha, InsR = Insulin Receptor, PD-L1 = Programmed cell death 1 ligand 1, TrkA = Tropomyosin receptor kinase A.

| Sequence Design Method | HA | IL7Rα | IR | PD-L1 | TrkA |
| --- | --- | --- | --- | --- | --- |
| **ProteinMPNN(Vanilla)** | 3.9% | 6.0% | 21.3% | 30.6% | 5.4% |
| **ProteinMPNN(Vanilla) + FastRelax** | **7.0%** | **11.1%** | 26.4% | 47.5% | 12.1% |
| **ProteinMPNN(Soluble)** | 5.0% | 5.8% | 27.6% | 33.9% | 8.5% |
| **ProteinMPNN(Soluble) + FastRelax** | 6.1% | 9.1% | **29.6%** | **52.3%** | **18.1%** |
| **FAMPNN** | 0.1% | 10.9% | 11.6% | 30.9% | 7.5% |
| **FAMPNN (target side-chains fixed)** | 1.6% | 4.8% | 8.0% | 9.9% | 4.1% |

We have extended the functionality of Full-Atom MPNN and Boltz-2 for binder design with additional metrics and parameters. Although Full-Atom MPNN extends ProteinMPNN and performs additional modelling of side-chains during sequence design, there was not a consistent improvement to success rates (Table 1). However, for some targets, it outperformed ProteinMPNN, highlighting the benefits of comparing sequence design software for different target classes.

Boltz-2 was more efficient than AlphaFold2 in generating structure predictions, but experimental testing is needed to compare their effectiveness in identifying promising designs. In a recent preprint containing a retrospective analysis of previous binder design campaigns^29^, a previous version of Boltz (Boltz-1) performed similarly to AlphaFold2 in identifying successful binders when using metrics derived from the prediction aligned error of interface residues. This study also indicates that when prediction scores are combined with biophysical metrics such as shape complementary and buried surface area they can be powerful selectors for successful designs. These metrics are captured in our analysis pipeline and can be leveraged to rank output designs.

Our protein design pipeline is constructed with several core design principles in mind. In designing ProteinDJ we aimed to address the current lack of standardised file formats and efficient data management, and establish consistent format conventions for protein design tools, structures and metadata. ProteinDJ assumes a linear design process i.e. sequential design steps without recycling of designs back through the pipeline. An automated iterative pipeline could be useful for some design cases to refine sequences and folds, if the pipeline avoids combinatorial explosion and can perform effective refinement unsupervised. However, we suggest that it would be preferable to intermittently inspect cycle outputs before embarking on computationally expensive large-scale design campaign.

Currently, ProteinDJ pipeline performs amino acid design of a single protein chain. In the future, we aim to implement recent tools such as RFdiffusion2/3^31; 32^ and LigandMPNN^33^ that can also perform protein design with small molecules. Although RFdiffusion can perform design of symmetric oligomers, we found that ProteinMPNN and FAMPNN were unable to perform symmetric sequence design without significant reprogramming, and so symmetric design was not enabled. However, as ProteinDJ was designed with modularity in mind, we envision future versions of this pipeline that include new and promising technologies.

Multiple pipelines for protein design have emerged recently with similarity to ProteinDJ. The BinderFlow pipeline^34^ consists of RFdiffusion, ProteinMPNN and AlphaFold2 Initial-Guess and includes a web dashboard for visualisation of results. BinderFlow also includes filtering and analysis steps between design stages but can only perform de novo binder design and partial diffusion of binders. It also requires manual installation of dependencies and relies on a SBATCH script, as opposed to containers and a flexible scheduling language as utilised by Nextflow. The code for RFdiffusion2^31^ was released during submission of our manuscript and included all-atom design modes, container support and integration of LigandMPNN and Chai-1 downstream, with example python scripts to run a pipeline. However, within a few months it was superseded by RFdiffusion3^32^, which integrates LigandMPNN for sequence design and RosettaFold3 for structure prediction in a different python-based framework (AtomWorks)^35^. BoltzGen^11^ uses a python-interface with conda packages or containers, to generate binders using a new generative model (BoltzGen), followed by inverted folding of designs for sequence generation (Boltz-IF) and structure prediction with Boltz-2, with filtering at the end of the pipeline to select designs. Additionally, while this manuscript was under review, a Nextflow-based pipeline called Ovo was released containing RFdiffusion and BindCraft, with support for community plugins including ProteinDJ^36^. Ovo is fully containerised with a web-interface for input specification and visualisation of metrics generated by an analytical ‘ProteinQC’ module and includes additional tools such as LigandMPNN and ColabDesign^37^ with improved support for multi-chain complexes. The shared Nextflow architecture and open-source nature of ProteinDJ and Ovo, has enabled integration of ProteinDJ into the Ovo pipeline.

In conclusion, our goal with this pipeline was two-fold: first, to package and implement protein design tools in a HPC environment at maximal efficiency; and secondly, to reduce the prohibitive complexity of these tools by providing an accessible interface with curated defaults and guidance for different design tasks. By incorporating different tools for sequence design and structure prediction, we allow for easy exploration and comparison of software tools for a design problem. Our open-source framework in ProteinDJ enables collaborative efforts to build similar workflows and incorporate future tools in this rapidly growing field of protein design.

## Materials and Methods

### Construction of ProteinDJ pipeline

We containerised all pipeline dependencies using Apptainer/Singularity^22^, summarized in Supplementary Table 1. The container images are available on GitHub, using the GitHub Container Repository (https://github.com/orgs/PapenfussLab/packages). ProteinDJ Nextflow profiles are set to pull the images for deployment on different HPC systems. Integration and containerisation required modification to the source code of some software packages as described below. Bindsweeper utilises Python3 packages including Click, PyYAML, and Pandas that requires installation of a python environment locally.

Generative AI coding tools (ChatGPT, Claude) were used to review and augment scripts and assist with troubleshooting. All outputs were critically reviewed, tested, and adapted by the authors. These tools were not used for writing or editing of the manuscript or for data analysis.

### Nextflow pipeline

We built the Nextflow pipeline for ProteinDJ with four main stages described in Figure 1 and implemented several high level parameters that control the execution mode and what programs are utilised for sequence design and structure prediction (Table 2). We also created parameters to allow skipping of stages or to run fold design only and exit, that are useful for testing or running comparisons e.g. structure prediction tools on the same set of binder designs.

**Table 2.** Parameters for controlling Nextflow workflow.

| Parameter | Default | Description |
| --- | --- | --- |
| design_mode | null | Pipeline mode. Options: monomer_denovo, monomer_foldcond, monomer_motifscaff, monomer_partialdiff, bindcraft_denovo, binder_denovo, binder_foldcond, binder_motifscaff, or binder_partialdiff |
| out_dir | ./pdj_results | Output directory for results. Existing results will be overwritten |
| seq_method | 'mpnn' | Sequence design method. Options: 'mpnn' (ProteinMPNN FastRelax) or 'fampnn' (Full-Atom MPNN). |
| pred_method | 'af2' | Structure prediction method. Options: 'af2' (AlphaFold2 Initial Guess) or 'boltz' (Boltz-2). |
| zip_pdbs | true | Whether to compress output final designs in a tar.gz archive. |
| rank_designs | true | Whether to rank final designs by prediction quality metrics. Ranking metric is automatically selected: af2_pae_interaction for AF2 or boltz_ptm_interface for Boltz-2. |
| max_designs | null | Maximum number of top-ranked designs to output (e.g. 100). |
| max_seqs_per_fold | null | Maximum number of sequences per fold to keep after ranking (e.g. 3). |
| run_fold_only | false | Whether to run only fold design and skip sequence design, prediction, and analysis. |
| skip_fold | false | Skip fold design, run sequence design, prediction, and analysis only. Requires valid skip_input_dir containing PDB and JSON files with metadata. Binder design PDBs must have binder as chain A and target as chain B. |
| skip_fold_seq | false | Skip fold design and sequence design, run structure prediction and analysis only. Requires valid skip_input_dir containing PDB files. Binder design PDBs must have binder as chain A and target as chain B. |
| skip_fold_seq_pred | false | Skip fold design, sequence design, and prediction, run analysis only. Requires valid skip_input_dir containing PDB files. Binder design PDBs must have binder as chain A and target as chain B. |
| skip_input_dir | null | Directory path for files when skipping stages (e.g. './rfd_results'). |
| rfd_models | './models/rfd' | Path to the RFdiffusion model checkpoints |
| af2_models | './models/af2' | Path to the AlphaFold2 models |
| boltz_models | './models/boltz' | Path to the Boltz-2 models |
| gpu_model | 'A30' | GPU model requested (e.g. 'A30') |
| gpu_queue | null | GPU queue name or partition |
| gpus | 4 | Number of GPUs to request |
| cpus_per_gpu | 8 | Number of CPUs requested per GPU |
| cpus | 24 | Number of CPUs requested for CPU-only jobs |
| memory_gpu | '24GB' | Memory requested for GPU jobs |
| memory_cpu | '24GB' | Memory requested for CPU-only jobs |

### RFdiffusion

The RFdiffusion environment configuration file was modified to resolve issues with a deprecated mode of installing PyTorch and NumPy^38^, and we also fixed a bug for fold-conditioning. We implemented a Groovy script that both validates and converts input parameters from Nextflow to RFdiffusion (see parameter list in Table 3). This script allows for segmentation of RFdiffusion runs into different modes and prevents conflict of parameter choices. Helper functions provide a human-readable interpretation of contigs and check that hotspot residues are present in input residues ranges to flag issues before launching the job. We added a python script to automatically generate contigs if not provided that include all residues in the input PDB (in GenerateRFDContigs), and a script to generate fold conditioning definition files for the target (in GenerateRFDFoldCond). The number of designs (for inference.num_designs) is divided across GPUs evenly, rounding up if needed. The output TRB files containing metadata are ‘unpickled’ and converted to JSON files.

**Table 3.** Parameters for fold generation using RFdiffusion (rfd) and BindCraft (bc)

| Parameter | Example Value | Description |
| --- | --- | --- |
| design_length | 60-70 | Design length of binder or monomer. e.g. '60' or '60-70'. Pipeline will randomly sample different sizes between these values. |
| num_designs | 8 | Number of designs to generate using RFdiffusion or BindCraft. |
| input_pdb | './target.pdb' | Path to an input PDB file. |
| hotspot_residues | 'A56,A115,A123' | Specifies hotspot residues for binding. |
| rfd_contigs | '[100-100/0 B101-221]' | Specifies the residue ranges to be diffused. |
| rfd_scaffold_dir | './binderscaffolds' | Path to directory containing scaffold files. |
| rfd_mask_loops | false | Whether to ignore loop regions when applying secondary structure constraints from scaffolds (default=true). Recommended to set to false for binder design as the scaffold sets have enough diversity |
| rfd_inpaint_seq | '[A17-19/A143-145]' | Residues that will have their sequence masked when diffusing |
| rfd_length | 100-120 | Length of diffused chain when performing motif scaffolding |
| rfd_partial_diffusion_timesteps | 20 | Number of timesteps for partial diffusion. |
| rfd_ckpt_override | 'complex_base' | Override the diffusion model checkpoint used by RFdiffusion. Choose from: 'active_site', 'base', 'base_epoch8', 'complex_base', 'complex_beta', 'complex_fold_base', 'inpaint_seq', 'inpaint_seq_fold'. RFdiffusion will automatically select a checkpoint. Not recommended to override except for 'binder_denovo' mode i.e. 'complex_beta' |
| rfd_noise_scale | 0.5 | Scale of noise for RFdiffusion translations (C-alpha) and rotations (frames). 1 (default) is recommended for non-binder modes to increase diversity. 0-0.5 is recommended for binder modes to increase success rates. |
| rfd_extra_config | 'contigmap.inpaint_str=[B165-178]' | Additional configuration parameters to pass to RFdiffusion that are not externalised in Nextflow. |
| bc_chains | null | Optional specification of input PDB chains. |
| bc_design_protocol | 'default' | Which binder design protocol to run? "default" is recommended. "betasheet" promotes the design of more beta sheeted proteins, but requires more sampling. "peptide" is optimised for helical peptide binders. |
| bc_template_protocol | 'default' | What target template protocol to use? "default" allows for limited amount of flexibility. "flexible" allows for greater target flexibility on both sidechain and backbone level. |
| bc_omit_AAs | 'C' | Residue types to avoid during sequence design |
| bc_fix_interface_residues | true | Whether to preserve/fix the interface residues of the binder designed by BindCraft during the subsequent sequence design stage |
| bc_advanced_json | true | Path to a custom advanced settings json file. |

### BindCraft

BindCraft is a self-contained protein design pipeline, which includes its own sequence design and structure prediction steps. We disabled these downstream steps using an in-built parameter (enable_mpnn=false), halting the BindCraft process after binder hallucination and before sequence design, and passing these outputs to our sequence design and structure prediction processes. We added a python script to prepare the input JSON files for BindCraft from the Nextflow parameters and another to process the output PDB files and metadata. Since BindCraft was integrated into ProteinDJ after RFdiffusion, converted PBD files follow RFdiffusion conventions e.g. chain A for binder, chain B for target, and we record the interface residue numbers so that users can optionally preserve the sequence of these residues during sequence design.

### ProteinMPNN FastRelax

We modified the python environment of ProteinMPNN FastRelax to use conda-forge instead of the Anaconda defaults channel and removed PyTorch and NVIDIA dependencies as we are using CPU-only functionality as recommended by Bennet *et al*^4^. Additionally, we switched to NumPy v2 and used OpenBLAS (Basic Linear Algebra Subprograms) over Intel MKL (Math Kernel Library) to resolve intermittent crashes when using FastRelax. We used the ‘inpaint_seq’ array from RFdiffusion to identify residues to retain during sequence design and add ‘FIXED’ remarks to the input PDB files to ProteinMPNN. To extend the functionality of the FastRelax protocol, we created a modified version of the dl_interface_design python script (dl_interface_design_multi.py). We created parameters to control the maximum number of cycles and convergence criteria to end cycles early, the number of sequences to generate between each relax cycle, and optional relaxation of the final design (see Table 4). We also generated a FastRelax protocol for monomer designs. When generating multiple sequences between relax cycles, the sequence with the lowest (best) score is automatically selected. When writing output PDB files, we also output a JSON file containing the design file name, sequence, and score.

**Table 4.** Parameters for sequence design using ProteinMPNN (mpnn) and Full-Atom MPNN (fampnn).

| Parameter | Default Value | Description |
| --- | --- | --- |
| seqs_per_design | 8 | Number of sequences to generate per RFdiffusion design. |
| mpnn_omitAAs | 'CX' | A string of all residue types (one letter case-insensitive) to avoid for design. |
| mpnn_temperature | 0.1 | Temperature for sequence sampling. Higher values increase sequence diversity. Recommended to lower significantly for binders to improve success (e.g., 0.0001). |
| mpnn_checkpoint_type | vanilla | Checkpoint type to use for ProteinMPNN. Choose from 'vanilla', 'soluble' or 'hyper' |
| mpnn_checkpoint_model | v_48_020 | Checkpoint model to use for ProteinMPNN. Indicates noise level used during training for backbone atoms e.g. 'v_48_020' noised with 0.20Å Gaussian noise. Choose from 'v_48_002','v_48_010','v_48_020','v_48_030'. |
| mpnn_backbone_noise | 0 | Standard deviation of Gaussian noise to add to backbone atoms. 0 = no change to backbone, 0.1-0.3 = allows mild perturbation of backbone. |
| mpnn_relax_max_cycles (previously relax_cycles) | 0 | Maximum number of fast relax cycles to run on each sequence. Set to zero to disable. |
| mpnn_relax_seqs_per_cycle (new) | 5 | Number of sequences to generate between relaxation steps. The sequence with the best score will be kept. |
| mpnn_relax_output (new) | false | Whether to run a relaxation cycle before saving output. |
| mpnn_relax_convergence_rmsd (new) | 0.2 | Design is considered converged if the C-alpha RMSD (Å) between cycles is <= this threshold. |
| mpnn_relax_convergence_score (new) | 0.1 | Design is considered converged if the improvement in score between cycles is <= this threshold. |
| mpnn_relax_convergence_max_cycles (new) | 1 | Design is considered converged if it meets both convergence criteria for n consecutive cycles. |
| mpnn_extra_config | null | Additional configuration parameters to pass to ProteinMPNN-FastRelax that are not externalised in Nextflow. |
| fampnn_fix_target_sidechains | false | Fix target side-chain positions when performing binder sequence design |
| fampnn_psce_threshold | 0.3 | Only keep sidechains below this PSCE threshold during design; null to keep all. |
| fampnn_temperature | 0.1 | Temperature for sequence sampling. Higher values increase sequence diversity. |
| fampnn_exclude_cys (new) | True | Exclude cysteine residues from sequence design. |
| fampnn_extra_config | null | Additional configuration parameters to pass to Full-Atom MPNN that are not externalised in Nextflow. |

**Table 5.**
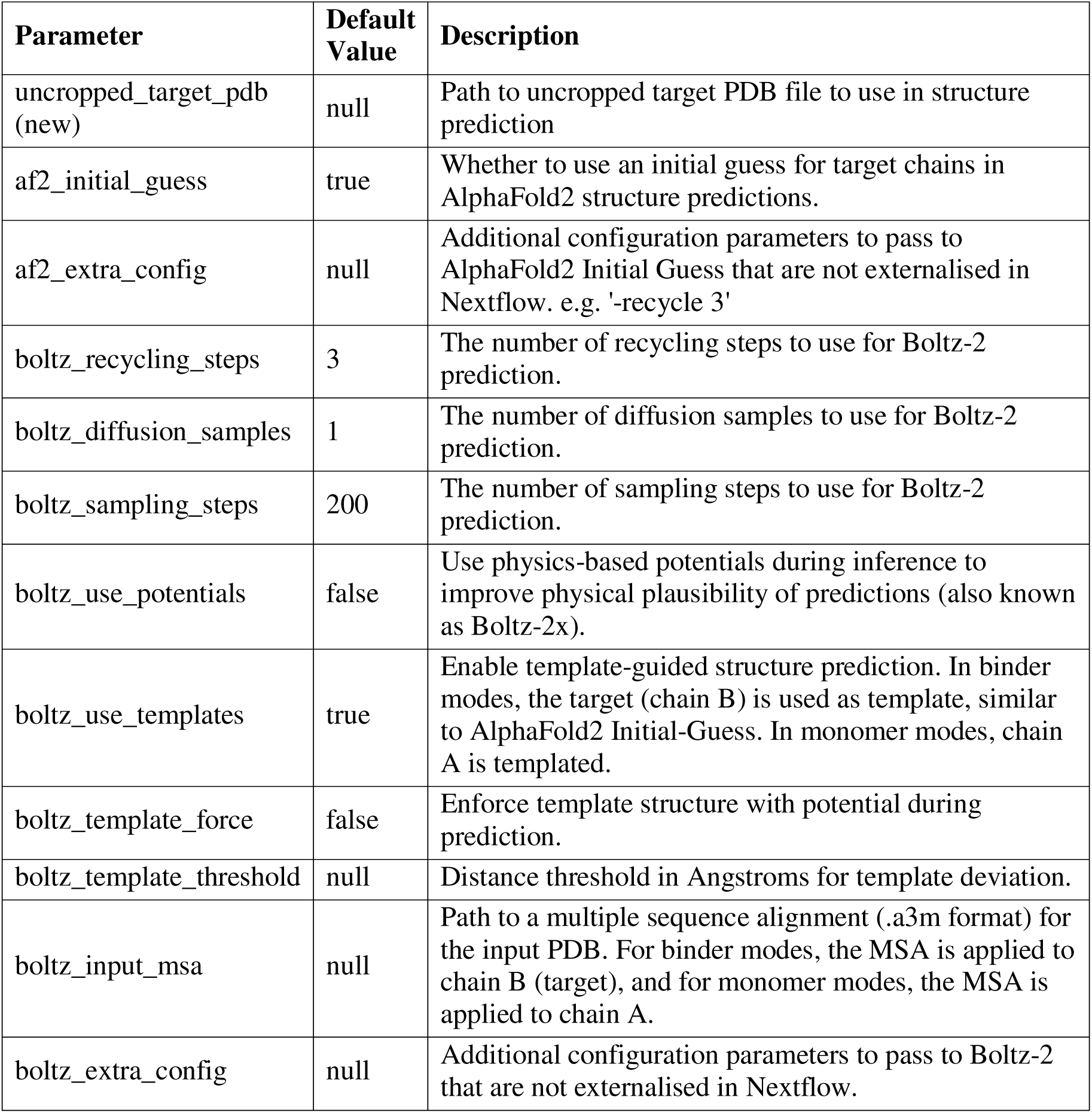
Parameters for structure prediction using AlphaFold2 Initial Guess (af2) and Boltz-2 (boltz)

### Full-Atom MPNN (FAMPNN)

Integration of FAMPNN required fixes to weight loading on GPUs. We modified the source code to add a parameter to exclude cysteine residues when performing sequence design (fampnn_exclude_cys). As with ProteinMPNN, we map fixed residues from RFdiffusion and exclude these from sequence design in the input CSV for FAMPNN. Since RFdiffusion removes all side-chain atoms from target residues, and FAMPNN requires sidechains for design context, we use PyRosetta to add sidechain atoms to fixed residues before design. Additionally, we derived a summary score for sequence design using the predicted side-chain error (PSCE) assigned to each atom by FAMPNN. This average PSCE is calculated per residue but excludes backbone atoms (that have a PSCE=0) and optionally C-beta atoms (that consistently have a PSCE ∼0.6), thereby weighting the score towards designed side-chain atoms. This score and the designed sequence are then output to a JSON metadata file.

### AlphaFold2 Initial Guess

For structure validation, we use the AlphaFold2 Initial Guess implementation from the dl-binder-design repository (https://github.com/nrbennet/dl_binder_design)^4^. Prior to containerisation we updated the deprecated PyRosetta installation method and resolved a deprecated import in the MMCIF parsing module that was causing install to fail. We observed that when the sequence length changed, AlphaFold2 generated new features so to improve efficiency we sort binders by size when batching to GPUs. As RFdiffusion scales poorly with target size for binder design tasks it is common practice to truncate target structures before design, but this can lead to issues in structure prediction, where the context is missing. We implemented new functionality to optionally replace the target chain in the protein designs with an uncropped target structure before structure prediction. The metrics from AlphaFold2 are converted to a JSON format, with binder_rmsd and target_rmsd renamed to rmsd_binder_bndaln and rmsd_binder_tgtaln for clarity. We added a rmsd calculation for all chains (rmsd_overall) and the target chain only (rmsd_target), as well as reporting of the predicted aligned error for all chains (pae_overall).

### Boltz-2

As Boltz-2 is a general purpose structure prediction program and not designed exclusively for protein design like AlphaFold2 Initial Guess, we constructed python scripts to generate input YAML files with target and design sequences (PrepBoltz). To speed up structure prediction, we disable generation of multiple sequence alignments (MSAs) and perform inference only, although the user can provide a MSA file for the target chain. We also provide the option of using the input target structure as templates to guide structure prediction, allowing Boltz-2 to function similarly to AlphaFold2 Initial-Guess. The metrics from Boltz2 are converted to a JSON format, with ‘ipde’, ‘iplddt’, and ‘iptm’ renamed to ‘pde_interface’, ‘plddt_interface’, and ‘ptm_interface’, respectively, for clarity. In the next process, AnalyseBoltz, we use scripts from https://github.com/DigBioLab/de_novo_binder_scoring to calculate additional structure prediction metrics based on the predicted aligned error (PAE) matrices, including: the PAE interaction, the minimum interaction prediction Score from Aligned Errors (ipSAE_min)^17; 29^, Local Interaction Score (LIS)^16^, and minimum predicted DockQ Score version 2 (pDockQ2_min)^15^. We also created a process, AlignBoltz, to align predictions with input designs to calculate RMSD values after alignment: overall RMSD of all C-alpha atoms, and for binder design, RMSD between the binder design and prediction, and RMSD between the target chains and prediction.

### Generation of scaffold templates for fold conditioning of binders

To assist users with performing fold conditioning of binders, we compiled structural templates of minibinders from published datasets^30; 39; 40^ and generated ∼23,000 PyTorch files directly compatible with RFdiffusion using a parallelised version of the ‘make_secstruc_adj.py’ script from the RFdiffusion GitHub (https://github.com/RosettaCommons/RFdiffusion) (Figure 4, Supplementary Table 5).

### Benchmarking and efficiency

We evaluated parallel scaling efficiency of the ProteinDJ pipeline using either 1, 2, 4, and 8 NVIDIA A30 GPUs on our SLURM HPC system. All executions used the maximum amount of GPUs and RAM and used the benchmarking target PD-L1 (Supplementary Table 6) generating 500 designs and 8 sequences per design. Scaling metrics were extracted from Nextflow trace files, with runtime calculated from actual compute time (start to completion timestamps) excluding queue time. Parallel efficiency was calculated as (T_1_ / (N × T_n_)) × 100%, where T_1_ = single-GPU runtime, N = GPU count, and T_n_ = N-GPU runtime. RFdiffusion was used for fold design, ProteinMPNN as the sequence design method and AF2 was used for structure prediction and validation.

To compare the efficiency of each software package, we generated 100 folds of *de novo* binders against PD-L1 with a sequence length of 50-100 residues using RFdiffusion and provided these to either ProteinMPNN, with and without a FastRelax cycle, or Full-Atom MPNN, generating 10 sequences per fold. All sequences were passed to either AlphaFold2 or Boltz-2 for prediction. ProteinMPNN was executed on nodes with 2.60GHz Intel Xeon E5-2690 CPUs and FAMPNN, AlphaFold2 and Boltz-2 utilised 4 NVIDIA A30 GPUs. We separately generated 100 folds (50-100 residues) with BindCraft using 4 NVIDIA A30 GPUs to compare the efficiency with RFdiffusion. The total runtime excluding queue time was divided by the number of designs.

### Comparison of sequence design methods

RFdiffusion in ProteinDJ was used to generate 100 folds for each target protein using previously described hotspots and contigs (Supplementary Table 6, Supplementary File 1). The noise scale for diffusion was set to zero for all runs. Using Bindsweeper, the folds were provided to ProteinMPNN or FAMPNN, followed by AlphaFold2 Initial Guess via the skip_input_dir path in ProteinDJ. The sequence design temperature that was set to 0.0001 in ProteinMPNN and FAMPNN for consistency. When using FastRelax, only one cycle was performed with one sequence per cycle. A design was considered successful if it passed three filters after prediction using AlphaFold2 Initial Guess: af2_max_rmsd_binder_bndaln ≤ 1, af2_max_pae_interaction ≤ 10, and af2_min_plddt_total ≥ 80 (see Supplementary Table 2 for definitions).

## Availability of code and data

ProteinDJ code is available at https://github.com/PapenfussLab/proteindj. An overview of all software dependencies and availability can be found in Supplementary Table 1. Benchmarking data can be found in Supplementary File 1.

## Acknowledgments

Development and evaluation of software was performed on the Walter and Eliza Hall Institute (WEHI) Milton HPC infrastructure. We would like to acknowledge the support and assistances of the WEHI Research Computing Platform. D.S. was supported by a WEHI Alan W Harris Honours Scholarship. J.M.H. was supported by an Australian National Health and Medical Research Council (NHMRC) EL1 Investigator Grant (2008096). A.T.P. was supported by an NHMRC Investigator Grant (2026643) and funding from the Lorenzo and Pamela Galli Medical Research Trust. This work was made possible through Victorian State Government Operational Support Program and the Australian Government NHMRC Independent Research Institutes Infrastructure Support Scheme to WEHI. A.P.T, I.S.L, and J.M.H. are members of the Australian Research Council Industrial Transformation Training Centre for Cryo-Electron Microscopy of Membrane Proteins for Drug Discovery (IC200100052). The contents of this published material are solely the responsibility of the individual authors and do not reflect the views of the NHMRC or funding partners.

## Author contributions

Dylan Silke: Conceptualization (equal); investigation (supporting); software (equal); visualization (supporting); writing – original draft (equal); writing – review and editing (equal).

Julie Iskander: Software (supporting); writing - review and editing (equal).

Junqi Pan: Software (supporting); visualization (supporting); writing - review and editing (equal).

Andrew P. Thompson; Supervision (supporting); writing - review and editing (equal).

Anthony T. Papenfuss; Supervision (equal); funding acquisition (equal); writing - review and editing (equal).

Isabelle S. Lucet; Supervision (equal); funding acquisition (equal); writing - review and editing (equal).

Joshua M. Hardy; Conceptualization (equal); investigation (lead); software (equal); supervision (equal); visualization (lead); writing – original draft (equal); writing - review and editing (equal).

## Competing interests

The authors declare no conflicts of interest.

---

**Supplementary Figure 1.**
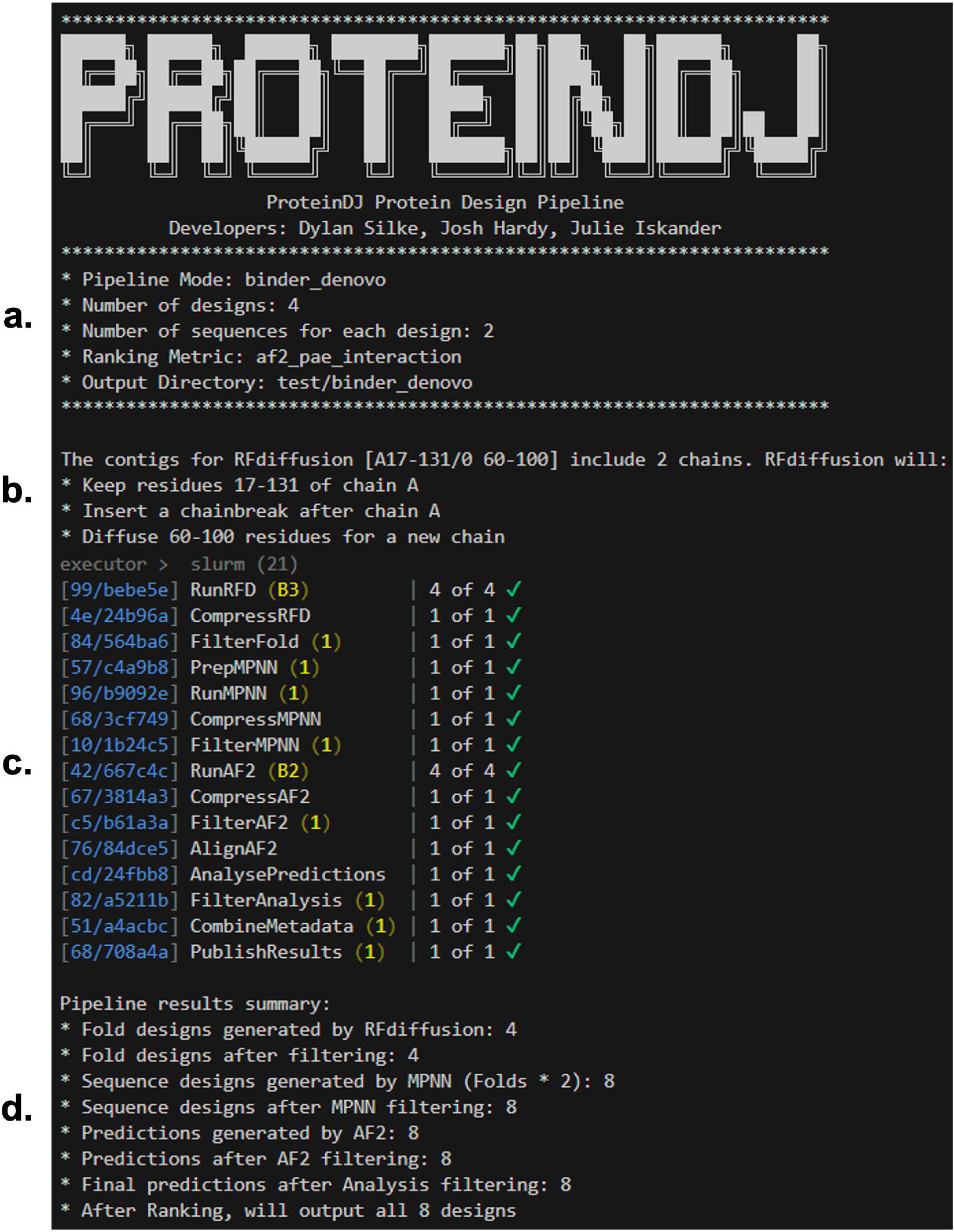
Example execution of ProteinDJ pipeline for *de novo* binder design. **a.** Header with key parameters and output directory. **b.** Description of contigs provided and how these will be used in diffusion. **c.** Standard Nextflow process tracker that updates as each task completes. **d.** A summary of the run, highlighting success rates at each stage.

**Supplementary Figure 2.**
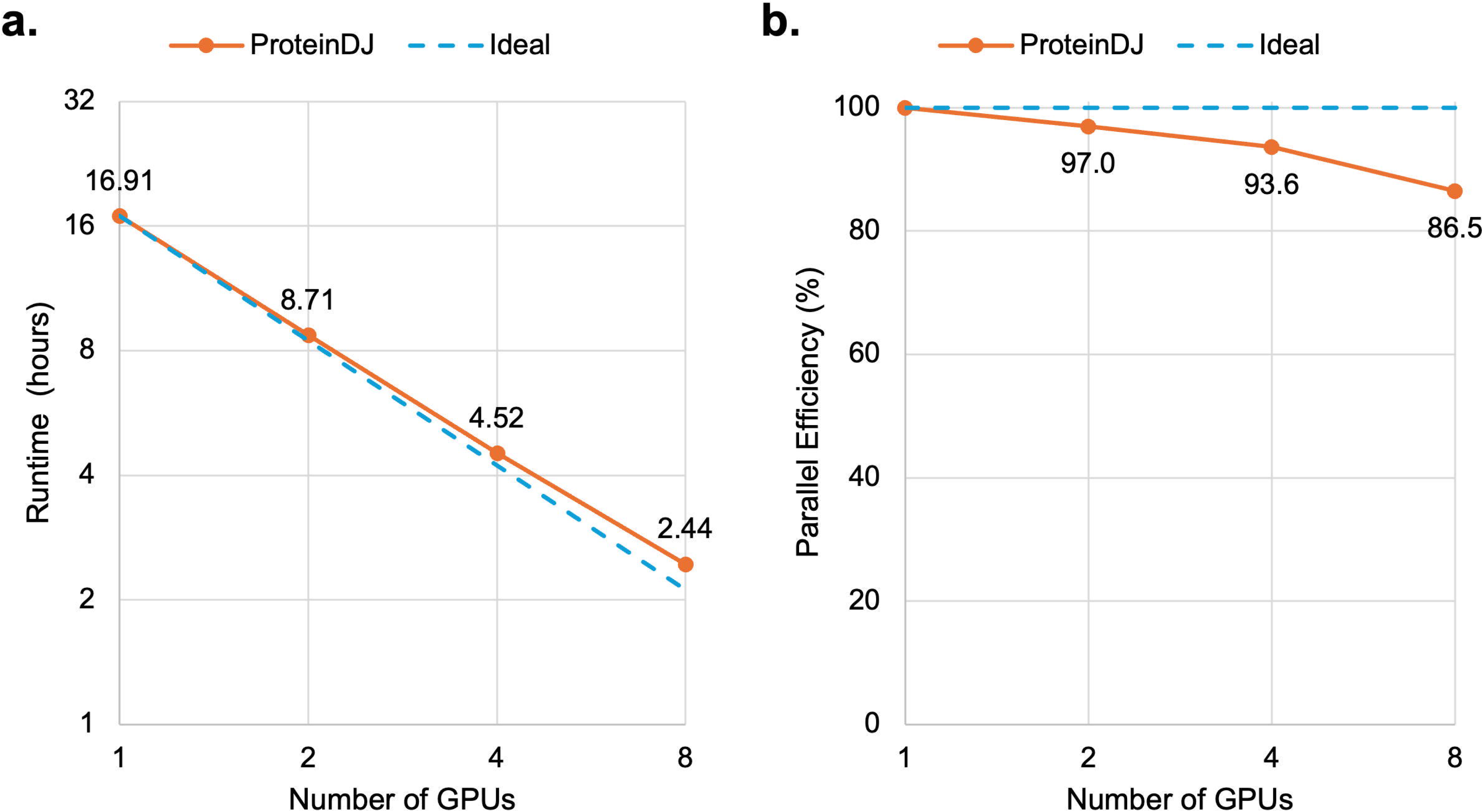
ProteinDJ multi-GPU efficiency and scaling. **(a)** Each data point represents the total wall-clock time excluding queue time for a complete ProteinDJ pipeline execution (4,000 designs) using the specified number of NVIDIA A30 GPUs. (b) Parallel efficiency of ProteinDJ calculated relative to single GPU execution. Each data point represents the parallel efficiency percentage calculated from wall-clock execution times.

**Supplementary Table 1.** The containers and software dependencies of ProteinDJ.

| Container | Description | Repository |
| --- | --- | --- |
| af2 | A modified version of AlphaFold2 (Initial Guess) as implemented in dl_binder_design. Used for structure prediction and design validation. | https://github.com/PapenfussLab/dl_binder_design.git |
| bindcraft | Contains the BindCraft binder design pipeline that includes AlphaFold2, ColabFold, ProteinMPNN and PyRosetta. | https://github.com/martinpacesa/BindCraft |
| boltz2 | Structure prediction design validation and filtering using Boltz-2 (inference only). | https://github.com/jwohlwend/boltz.git |
| dl_binder_design | Environment for ProteinMPNN FastRelax used for sequence design. Includes ‘vanilla’ and ‘soluble’ checkpoint models and implementation of Rosetta FastRelax Protocol | https://github.com/PapenfussLab/dl_binder_design.git |
| fampnn | Environment for Full-Atom MPNN, a sequence design tool. | https://github.com/PapenfussLab/fampnn.git |
| pyrosetta_tools | Python interface to Rosetta molecular modeling suite (PyRosetta) bundled with standard data analysis libraries (BioPython, NumPy, Pandas, Matplotlib) for processing pipeline results. | N/A |
| python_tools | Lightweight container with standard python data analysis libraries (BioPython, NumPy, Pandas, Matplotlib) for processing pipeline results. | N/A |
| rfdiffusion | Contains RFdiffusion, a fold design software integral to the pipeline | https://github.com/PapenfussLab/RFdiffusion |

**Supplementary Table 2.** The filtering parameters available in ProteinDJ.

| **Parameter** | **Description** |
| --- | --- |
| **RFdiffusion** |  |
| fold_min_helices | Minimum number of alpha-helices required. |
| fold_max_helices | Maximum number of alpha-helices allowed. |
| fold_min_strands | Minimum number of beta-strands required. |
| fold_max_strands | Maximum number of beta-strands allowed. |
| fold_min_ss | Minimum number of secondary structure elements (α-helices + β-strands). |
| fold_max_ss | Maximum number of secondary structure elements (α-helices + β-strands). |
| fold_min_rog | Minimum radius of gyration (Å). |
| fold_max_rog | Maximum radius of gyration (Å). |
| **ProteinMPNN** |  |
| mpnn_max_score | Maximum MPNN score (negative log likelihood). |
| **FAMPNN** |  |
| fampnn_max_psce | Max predicted-side chain error (PSCE) score for designed side-chains. |
| **AlphaFold2 Initial Guess** |  |
| af2_max_pae_interaction | Max predicted aligned error for interactions |
| af2_max_pae_overall | Max predicted aligned error for all chains |
| af2_max_pae_binder | Max predicted aligned error for binder |
| af2_max_pae_target | Max predicted aligned error for target |
| af2_min_plddt_overall | Max C-alpha RMSD of binder when binder chains are aligned |
| af2_min_plddt_binder | Max C-alpha RMSD of binder when target chains are aligned |
| af2_min_plddt_target | Max C-alpha RMSD of target when target chains are aligned |
| af2_max_rmsd_overall | Max C-alpha RMSD when all chains are aligned |
| af2_max_rmsd_binder_bndaln | Max C-alpha RMSD of binder when binder chains are aligned |
| af2_max_rmsd_binder_tgtaln | Max C-alpha RMSD of binder when target chains are aligned |
| af2_max_rmsd_target | Max C-alpha RMSD of target when target chains are aligned |
| **Boltz-2** |  |
| boltz_max_rmsd_overall | Max C-alpha RMSD between all chains of Boltz-2 prediction and RFD design |
| boltz_max_rmsd_binder | Max C-alpha RMSD between binder chains of Boltz-2 prediction and RFD design. |
| boltz_max_rmsd_target | Max C-alpha RMSD between target chains of Boltz-2 |
|  | prediction and RFD design. |
| boltz_min_conf_score | Minimum confidence score of the prediction |
| boltz_min_ptm | Minimum predicted template modelling (pTM) score of the prediction |
| boltz_min_ptm_interface | Minimum pTM-score of the prediction interface |
| boltz_min_ptm_binder | Minimum pTM-score of the binder chain |
| boltz_min_ptm_target | Minimum pTM-score of the target chain |
| boltz_min_plddt | Minimum pLDDT score of the prediction |
| boltz_min_plddt_interface | Minimum pLDDT score of the prediction interface |
| boltz_max_pde | Maximum predicted distance error of the prediction |
| boltz_max_pde_interface | Maximum predicted distance error of the prediction interface |
| boltz_min_ipSAE_min | Minimum value allowed for the minimum interaction prediction Score from Aligned Errors (ipSAE) of target and binder chains |
| boltz_min_LIS | Minimum Local Interaction Score (LIS) |
| boltz_min_pDockQ2_min | Minimum value allowed for the minimum predicted DockQ Score v2 of target and binder chains. |
| boltz_max_pae_interaction | Maximum predicted aligned error at interaction interfaces |
| **PyRosetta/BioPython** |  |
| pr_min_helices | Minimum number of alpha-helices in predicted structure |
| pr_max_helices | Maximum number of alpha-helices in predicted structure |
| pr_min_strands | Minimum number of beta-strands in predicted structure |
| pr_max_strands | Maximum number of beta-strands in predicted structure |
| pr_min_total_ss | Minimum total secondary structure elements (α-helices + β-strands) in predicted structure |
| pr_max_total_ss | Maximum total secondary structure elements (α-helices + β-strands) in predicted structure |
| pr_min_rog | Minimum radius of gyration ($\text{\AA}$) of predicted structure |
| pr_max_rog | Maximum radius of gyration ($\text{\AA}$) of predicted structure |
| pr_min_intface_bsa | Minimum buried surface area ($\text{\AA}^2$) at the binding interface |
| pr_min_intface_shpcomp | Minimum shape complementarity of interface (0-1 scale; 1 is optimal) |
| pr_min_intface_hbonds | Minimum number of hydrogen bonds at the interface |
| pr_max_intface_unsat_hbonds | Maximum number of buried, unsatisfied hydrogen bonds at the interface |
| pr_max_intface_deltag | Maximum solvation free energy gain at interface (Rosetta Energy Units; lower is better) |
| pr_max_intface_deltagtobsa | Maximum ratio of delta-G to buried surface area |
| pr_min_intface_packstat | Minimum packing quality of the interface (0-1 scale; higher is better) |
| pr_max_tem | Maximum total energy metric score (Rosetta Energy |
|  | Units; lower indicates more stable designs) |
| pr_max_surfhphobics | Maximum percentage of hydrophobic residues exposed on the surface |
| pr_max_sap | Maximum mean residue Spatial Aggregation Propensity for monomer/binder (solubility prediction; lower is better) |
| pr_max_sap_complex | Maximum mean residue Spatial Aggregation Propensity for binder in complex (solubility prediction; lower is better) |
| seq_min_ext_coef | Minimum extinction coefficient at 280nm ($M^{-1}cm^{-1}$) |
| seq_max_ext_coef | Maximum extinction coefficient at 280nm ($M^{-1}cm^{-1}$) |
| seq_min_pi | Minimum isoelectric point (pI) of the sequence |
| seq_max_pi | Maximum isoelectric point (pI) of the sequence |

**Supplementary Table 3.** ProteinDJ outputs a wide array of additional metrics to assist researchers to evaluate potential designs. Some metrics are calculated by software in the pipeline (^+^), but there are additional metrics calculated using PyRosetta^18^ and BioPython^19^.

| Metric name | Explanation |
| --- | --- |
| description | Unique filename of structure prediction |
| fold_id | Fold ID for independent execution of RFdiffusion |
| seq_id | Sequence ID for independent execution of ProteinMPNN |
| rfd_sampled_mask<sup>+</sup> | Contigs used by RFdiffusion to produce this fold (e.g., ['E6-155/0', '100-100']) |
| bc_length | Length of binder selected by BindCraft |
| bc_plddt | Average per-residue confidence score (0-100) for the complex calculated by BindCraft. |
| bc_rmsd_target | The RMSD between the input target structure and the target structure after binder design |
| fold_helices | Number of α-helices in designed fold |
| fold_strands | Number of β-strands in designed fold |
| fold_total_ss | Total secondary structures (helices + strands) in designed fold |
| fold_RoG | Radius of gyration (RoG) for designed fold (Å) |
| rfd_time | Time (seconds) taken by RFdiffusion to produce a fold |
| bc_time | Time (seconds) taken by BindCraft to produce a fold |
| fampnn_avg_psce | Average predicted sidechain error (PSCE) from FAMPNN |
| mpnn_score<sup>+</sup> | ProteinMPNN's average negative log likelihood |
| af2_pae_interaction<sup>+</sup> | AF2 PAE for binding interface residue pairs (Å) |
| af2_pae_overall | AF2 PAE for all residue positions (Å) |
| af2_pae_binder<sup>+</sup> | AF2 predicted aligned error (PAE) for binder residue positions (Å) |
| af2_pae_target<sup>+</sup> | AF2 PAE for target protein residue positions (Å) |
| af2_plddt_overall<sup>+</sup> | Average predicted Local Distance Difference Test (pLDDT) score for all residues |
| af2_plddt_binder<sup>+</sup> | Average pLDDT score for binder residues |
| af2_plddt_target<sup>+</sup> | Average pLDDT score for target residues |
| af2_rmsd_overall | C-alpha RMSD comparing RFdiffusion design and AF2 prediction (all chains). |
| af2_rmsd_binder_tgtaln<sup>+</sup> | C-alpha RMSD between RFD binder and AF2 binder when target chain is aligned (Å) |
| af2_rmsd_binder_bndaln<sup>+</sup> | C-alpha RMSD between RFD binder and AF2 binder when binder chain is aligned (Å) |
| af2_rmsd_target | C-alpha RMSD between RFD target and AF2 target when binder chain is aligned (Å) |
| af2_time<sup>+</sup> | Time (seconds) for AlphaFold2 prediction |
| boltz_rmsd_overall | C-alpha RMSD between all chains of Boltz-2 prediction and RFD design (Å) |
| boltz_rmsd_binder | C-alpha RMSD between binder chains of Boltz-2 prediction |
|  | and RFD design (Å) |
| boltz_rmsd_target | C-alpha RMSD between target chains of Boltz-2 prediction and RFD design (Å) |
| boltz_conf_score<sup>+</sup> | Confidence score for Boltz-2 prediction |
| boltz_ipSAE_min | Minimum interaction Prediction Score from Aligned Errors (ipSAE) of target and binder chains |
| boltz_LIS | Local Interaction Score (LIS) |
| boltz_pDockQ2_min | Minimum predicted DockQ Score v2 (calculated from PAE) of target and binder chains. |
| boltz_pae_interaction | Predicted aligned error for interaction interfaces. |
| boltz_ptm<sup>+</sup> | The predicted template modelling (pTM) score for the Boltz-2 prediction |
| boltz_ptm_interface<sup>+</sup> | Interface pTM-score for Boltz-2 prediction |
| boltz_ptm_binder<sup>+</sup> | Predicted template modelling score of binder chain |
| boltz_ptm_target<sup>+</sup> | Predicted template modelling score for target chain |
| boltz_plddt<sup>+</sup> | pLDDT score for Boltz-2 prediction |
| boltz_plddt_interface<sup>+</sup> | Interface pLDDT score for Boltz-2 prediction of the complex |
| boltz_pde<sup>+</sup> | Predicted distance error (PDE) for Boltz-2 prediction |
| boltz_pde_interface<sup>+</sup> | Interface PDE for Boltz-2 prediction of the complex (Å) |
| pr_helices | Number of α-helices in predicted structure |
| pr_strands | Number of β-strands in predicted structure |
| pr_total_ss | Total secondary structures (helices + strands) in predicted structure |
| pr_RoG | RoG of predicted structure (Å) |
| pr_intface_BSA | Buried surface area (BSA) at binding interface (Å<sup>2</sup>) |
| pr_intface_shpcomp | Interface shape complementarity (0-1, 1=optimal) |
| pr_intface_hbonds | Number of interface hydrogen bonds |
| pr_intface_unsat_hbonds | Number of buried, unsatisfied hydrogen bonds at the interface. |
| pr_intface_deltaG | Solvation free energy gain (delta G) at interface (kcal/mol) |
| pr_intface_deltaGtoBSA | Ratio of delta-G to buried surface area |
| pr_intface_packstat | Interface packing quality (0-1, higher=better) |
| pr_TEM | Rosetta Total Energy Metric (TEM) score of design (lower=more stable) |
| pr_surfhphobics_% | Percentage of hydrophobic residues on design surface |
| pr_SAP | Mean residue Spatial Aggregation Propensity of monomer/binder (solubility prediction; lower is better). |
| pr_SAP_complex | Mean residue Spatial Aggregation Propensity of binder when complexed with target (solubility prediction; lower is better). |
| seq_ext_coef | Extinction coefficient (M<sup>-1</sup>cm<sup>-1</sup>) at 280nm |
| seq_length | Number of amino acids in design |
| seq_MW | Molecular weight (MW) of design (Da) |
| seq_pI | Isoelectric point (pI) of design |
| sequence | Amino acid sequence of design |

+Existing metrics from RFdiffusion, ProteinMPNN-Fast Relax, AlphaFold 2 Initial Guess and Boltz-2

**Supplementary Table 4.**
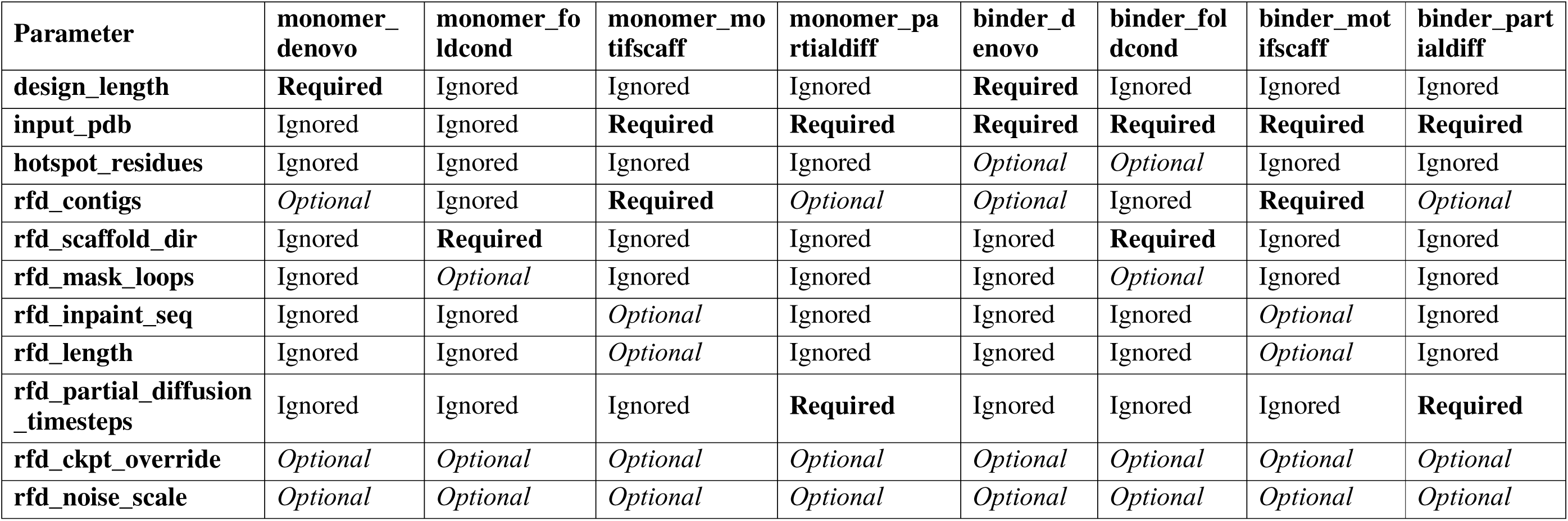
Parameters for RFdiffusion used for different modes. Parameters are ignored if they are not relevant to the design type or cause conflicts.

**Supplementary Table 5.** Scaffold templates for fold conditioning.

| **Prefix** | **Description** | **Count** |
| --- | --- | --- |
| HHH | "3-helical bundles" | 17668 |
| HHHH | "4-helical bundles" | 3499 |
| HEEHE | "Mixed alpha-helical (H) and beta-strand (E) scaffold. Artificial." | 208 |
| EHEEHE | "Mixed alpha-helical (H) and beta-strand (E) scaffold based on ferredoxin" | 1555 |

**Supplementary Table 6.** Benchmarking target structures. Structures were prepared including the same residues as used for RFdiffusion benchmarking in Watson et al. ^5^. The contigs and hotspots provided to RFdiffusion are indicated below.

| **Protein (Acronym)** | **Biological function** | **PDB ID** | **RFdiffusion contigs** | **RFdiffusion hotspots** |
| --- | --- | --- | --- | --- |
| Influenza A H1 haemagglutinin (HA) | Viral surface protein for host cell attachment and entry. | 5VLI | [A4-53/A79-83/A110-114/A261-325/0 B501-568/B580-670/0 50-100] | B521, B545, B552 |
| Interleukin-7 receptor alpha (IL-7Rα) | Receptor for IL-7, important for lymphocyte development. | 3DI3 | [B17-209/0 50-100] | B58, B80, B139 |
| Insulin Receptor (InsR) | Insulin receptor, regulates glucose uptake and metabolism. | 4ZXB | [E6-155/0 50-100] | E64, E88, E96 |
| Programmed cell death 1 ligand 1 (PD-L1) | Immune checkpoint, protein helps regulate immune responses. | 5O45 | [A17-131/0 50-100] | A56, A115, A123 |
| Tropomyosin receptor kinase A (TrkA) | Receptor for nerve growth factor, promotes neuron survival and growth. | 1WWW | [X282-382/0 50-100] | X294, X296, X333 |

## Notes

### Competing Interest Statement

The authors have declared no competing interest.

### Summary of Updates

Manuscript updated to reflect integration of BindCraft and updates to parameters and metrics in ProteinDJ v2; minor revisions to figures and main text; new section in Discussion comparing recent pipelines; Supplemental files updated.

